# Bridging the spatiotemporal scales of macromolecular transport in crowded biomimetic systems

**DOI:** 10.1101/431528

**Authors:** Kathryn Regan, Devynn Wulstein, Hannah Rasmussen, Ryan McGorty, Rae M. Robertson-Anderson

## Abstract

Crowding plays a key role in the transport and conformations of biological macromolecules. Gene therapy, viral infection and transfection require DNA to traverse the crowded cytoplasm, including a heterogeneous cytoskeleton of filamentous proteins. Given the complexity of cellular crowding, the dynamics of biological molecules can be highly dependent on the spatiotemporal scale probed. We present a powerful platform that spans molecular and cellular scales by coupling single-molecule conformational tracking (SMCT) and selective-plane illumination differential dynamic microscopy (SPIDDM). We elucidate the transport and conformational properties of large DNA, crowded by custom-designed networks of actin and microtubules, to link single-molecule conformations with ensemble DNA transport and cytoskeleton structure. We show that actin crowding leads to DNA compaction and suppression of fluctuations, combined with anomalous subdiffusion and heterogeneous transport, whereas microtubules have much more subdued impact across all scales. Interestingly, in composite networks of both filaments, microtubules primarily govern single-molecule DNA dynamics whereas actin governs ensemble transport.

## Introduction

The impact of crowding on macromolecular transport and conformations is a topic of intensive theoretical and experimental research.^1–9^ The primary inspiration for much of this work is to understand how large biomacromolecules such as nucleic acids and proteins can function in the highly crowded environment of the cell. For example, DNA transport through the crowded cytoplasm is essential for transformation, transcription, looping, gene expression, viral infection and gene therapy.^1,2,9,10^ Crowding is arguably the most important factor in determining passive intracellular transport properties, and also plays a key role in determining the conformations, and thus stability and function, of macromolecules.

Most in vitro studies aimed at understanding the role that crowding plays in macromolecular conformations or transport have focused on the effect of small soluble proteins, often mimicked with synthetic crowders such as dextran or PEG.^3,11–14^ Several of these studies have reported normal Brownian motion of spherical tracers and DNA, in which the mean-squared displacement in each direction (MSD) scales linearly with time (i.e. *MSD = 2Dt* where *D* is the diffusion coefficient).^15–18^ However, others have found anomalous subdiffusion where *MSD = Kt^α^* with *K* being the transport coefficient and *α* being in the range of 0.4–0.9.^4,19–21^ These studies have also shown evidence of macromolecular compaction, swelling, and elongation depending on type and size of tracer macromolecule and crowder.^4,5,22^ These crowding-induced conformational changes in turn impact the resulting transport properties.

Much less appreciated is the fact that the cytoplasm of eukaryotic cells is also crowded by pervasive networks of filamentous proteins that comprise the cytoskeleton.^23,24^ Given the size and rigidity of cytoskeletal proteins, the steric interactions between them, and the viscoelastic nature of the networks, the cytoskeleton has been identified as a key barrier to efficient gene delivery, and its impact on DNA transport is likely highly unique to that of the widely-studied viscous systems of soluble proteins.^4,15,19,25^ Two ubiquitous cytoskeletal proteins are semiflexible actin filaments, with a persistence length of ~10 μm and thickness of ~7 nm, and rigid microtubules, with a persistence length of ~1 mm and thickness of ~25 nm.^4,23,26,27^ At physiologically relevant concentrations, both filaments exhibit wide length distributions of ~1 – 50 μm and form sterically interacting networks with mesh sizes of ~0.1 – 1 μm^-1^). The resulting environment is thus highly heterogeneous on the lengthscale of diffusing macromolecules.

This inherent heterogeneity, similar to other biological networks such as mucus and the extracellular matrix, limits the ability of standard particle-tracking and scattering methods to individually gather a comprehensive picture of the complex macromolecular dynamics that occur over a wide range of scales. For example, measurements of nanospheres diffusing in actin networks have reported diffusion coefficients that vary over 2 orders of magnitude depending on the timescale probed.^28^ Similar studies have reported subdiffusion in actin networks at intermediate times but normal diffusion at short and long times.^29^ These results highlight the importance of measurements with fast time resolution *and* long timescales. This effect is amplified when examining environments composed of more than one crowder species, where the range of dynamics are even more spread out in time and space.^30^ Single-molecule tracking can quantify the distributions of conformational dynamics and trajectories of single diffusing macromolecules. However, these methods are limited by a need for a large ensemble of molecules to accurately determine MSDs and can often only measure transport over relatively short length and timescales due to photobleaching and diffusion beyond a limited region-of-interest. Conversely, scattering methods, which investigate bulk patterns in diffusion, can measure transport properties over large spatiotemporal scales, but they are unable to discern individual molecular trajectories or conformations.

Here we present a powerful platform that combines single molecule conformational tracking with digital Fourier microscopy to allow for the coupling of crowding-induced conformations and trajectories of single molecules with ensemble-averaged transport properties across a large range of length and time scales (~0.05 – 60 s, ~0.1 – 25 μm) (Fig. 1). This platform also allows for direct coupling of macromolecular dynamics to the structural properties and heterogeneities of the environment. The wide spatiotemporal range and variety of measurement deliverables is made possible by integrating two recently developed complementary techniques: selective-plane illumination differential dynamic microscopy (SPIDDM)^31^ and single-molecule conformational tracking (SMCT).^3,6,32^

**Figure 1.**
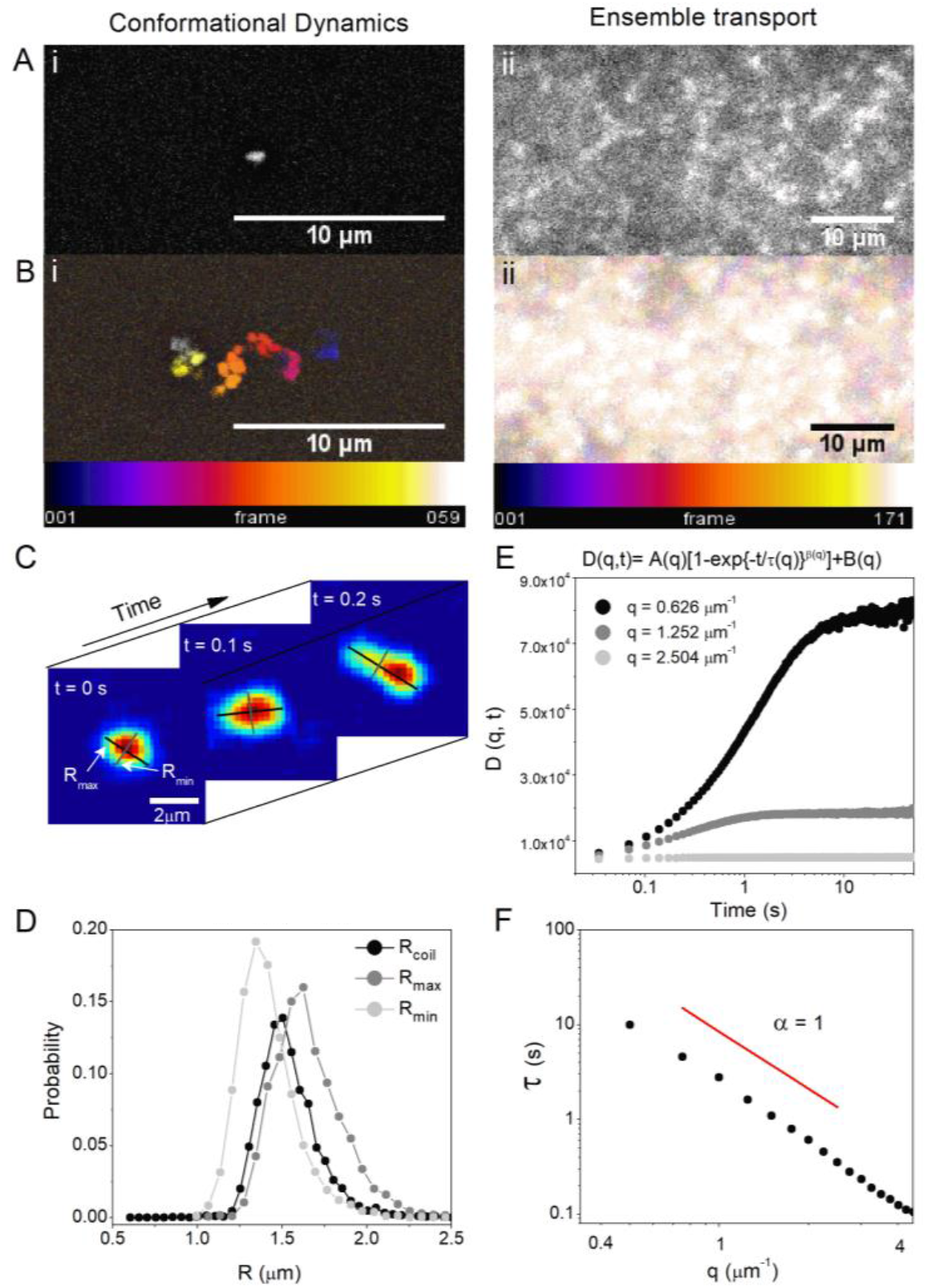
Coupling SMCT with SPIDDM enables robust measurements of macromolecular conformations and transport properties over a wide spatiotemporal range. (A) High-resolution epifluorescence imaging resolves conformations of single DNA molecules (i) while LSM imaging over large field-of-views captures ensemble DNA dynamics (ii). (B) Temporal color maps of videos of DNA acquired at 10 fps using epifluorescence (i) and 29 fps using LSM (ii). The combined method allows for resolution of single-molecule shape dynamics and trajectories (i) as well as statistically robust ensemble transport properties (ii). (C) SMCT measures major and minor axis lengths *R_max_* and *R_min_* of each molecule for each frame of each video. (D) Distributions of *R_max_* and *R_min_* for the entire ensemble of molecules and frames, as well as the distribution of coil sizes *Rcoil = [½(R_max^2^_+R_min^2^_)]^1/2^* quantifies the population of conformational sizes and shapes accessed by molecules. (E) DDM algorithms compute the radial average of Fourier transforms of differences of images separated by a given lag time. This analysis provides the image structure function *D(q,t)*, which is fit to a stretched exponential to extract decay times for each accessible spatial frequency. (F) To describe ensemble diffusion properties, the characteristic decay time *τ* for a given spatial frequency *q* is fit to a power-law relation *τ = (Kq^2^)^-1/α^* where *K* is the transport coefficient and *α* is the diffusive scaling exponent. All data shown is for DNA in buffer conditions (see text for details).

SPIDDM couples light-sheet microscopy (LSM), an optical sectioning microscopy technique that allows for minimum exposure to excitation light,^33–35^ with differential dynamic microscopy (DDM), a type of digital Fourier microscopy that quantifies intensity fluctuations within a time series of images.^36^ Importantly, DDM does not require that individual DNA molecules be localized, let alone tracked, in order to determine their transport dynamics. Therefore, larger fields-of-view and less excitation power can be used to extract transport dynamics as compared to single-molecule tracking methods.^31^ Complementing the efficient extraction of ensemble transport dynamics with DDM, we use SMCT to resolve the position, size and shape of individual molecules over time. These conformational measurements include the time-resolved distributions of major and minor axis lengths (*R_max_, R_min_*) and the spatiotemporal scales over which conformational fluctuations occur (Fig. 1).

Using this combined approach, we characterize the dynamics of large naked DNA molecules diffusing in custom-engineered cytoskeletal networks, allowing for direct coupling of DNA diffusivity and conformational dynamics with the properties of the surrounding heterogeneous protein environment. In order to model the cytoskeleton, we create tunable, well-characterized networks of actin filaments and microtubules, as well as a composite of both proteins (Fig 2).^30^ Ensemble analysis reveals that DNA transport in actin and composite networks is highly heterogeneous as well as subdiffusive, whereas microtubule networks induce far less heterogeneity and restricted diffusion. SMCT analysis reveals that actin networks appreciably compact DNA and suppress conformational fluctuations, while microtubule and composite networks result in much less compaction and less constrained fluctuations. These results demonstrate the integral role that coupling dynamics across a wide range of scales plays in characterizing macromolecular dynamics in crowded heterogeneous systems.

**Figure 2.**
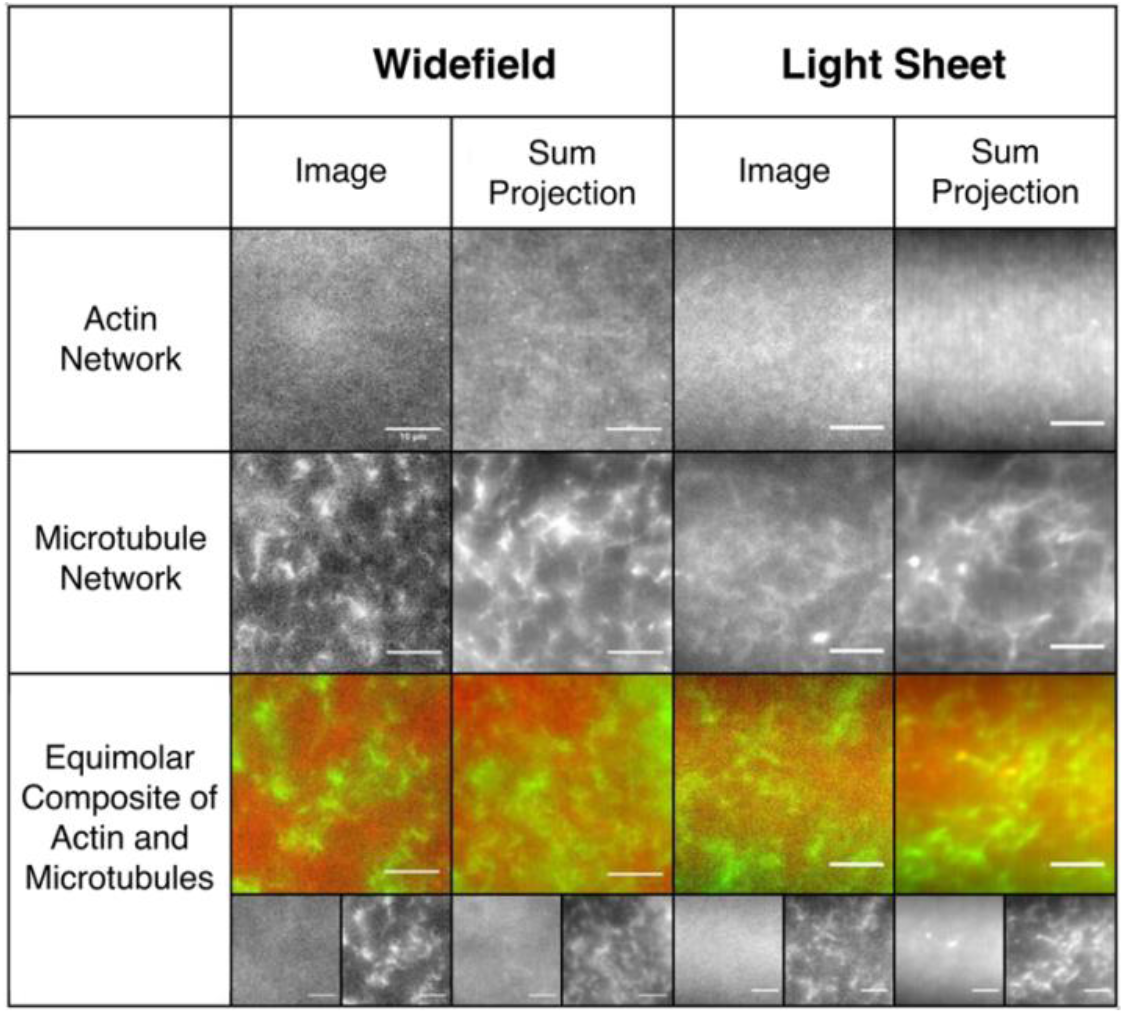
In vitro cytoskeleton networks exhibit robust characteristics and structure. Representative images (left) and sum projections of videos (right) taken using the epifluorescence and lightsheet microscopy methods used for SMCT and SPIDDM measurements. All scale bars represent 10 μm. As shown, networks formed in the respective sample chambers are nearly indistinguishable from one another. Actin networks have a smaller mesh size than microtubule networks, exhibited as more uniformity at the microscale. Microtubules form more rigid structures, exhibited as higher contrast projections compared to actin. In composites, actin and microtubules are well-mixed and sterically interacting with no large-scale phase separation. In all images, actin and tubulin are labeled with Alexa-488 (1:5 ratio labeled:unlabeled monomers) and rhodamine (1:35 labeled:unlabeled dimer ratio). Videos are all 400 frames acquired at 40 fps.

## Results and Discussion

We use SMCT to determine the effect of cytoskeleton network confinement on the conformational dynamics of single DNA molecules (Fig. 1, Methods).^3,6,32^ Specifically, we evaluate the distribution of shapes and sizes of molecular conformations and determine how the measured conformations change over time. The distributions of DNA coil sizes *R_coil_* = *[½(R_max^2^_ + R_min^2^_)]^1/2^* for each condition clearly show that all cytoskeleton networks induce some degree of DNA compaction compared to buffer conditions (Fig. 3A). The degree of compaction is highest for actin filaments while microtubules and composite networks induce comparably less compaction.

**Figure 3.**
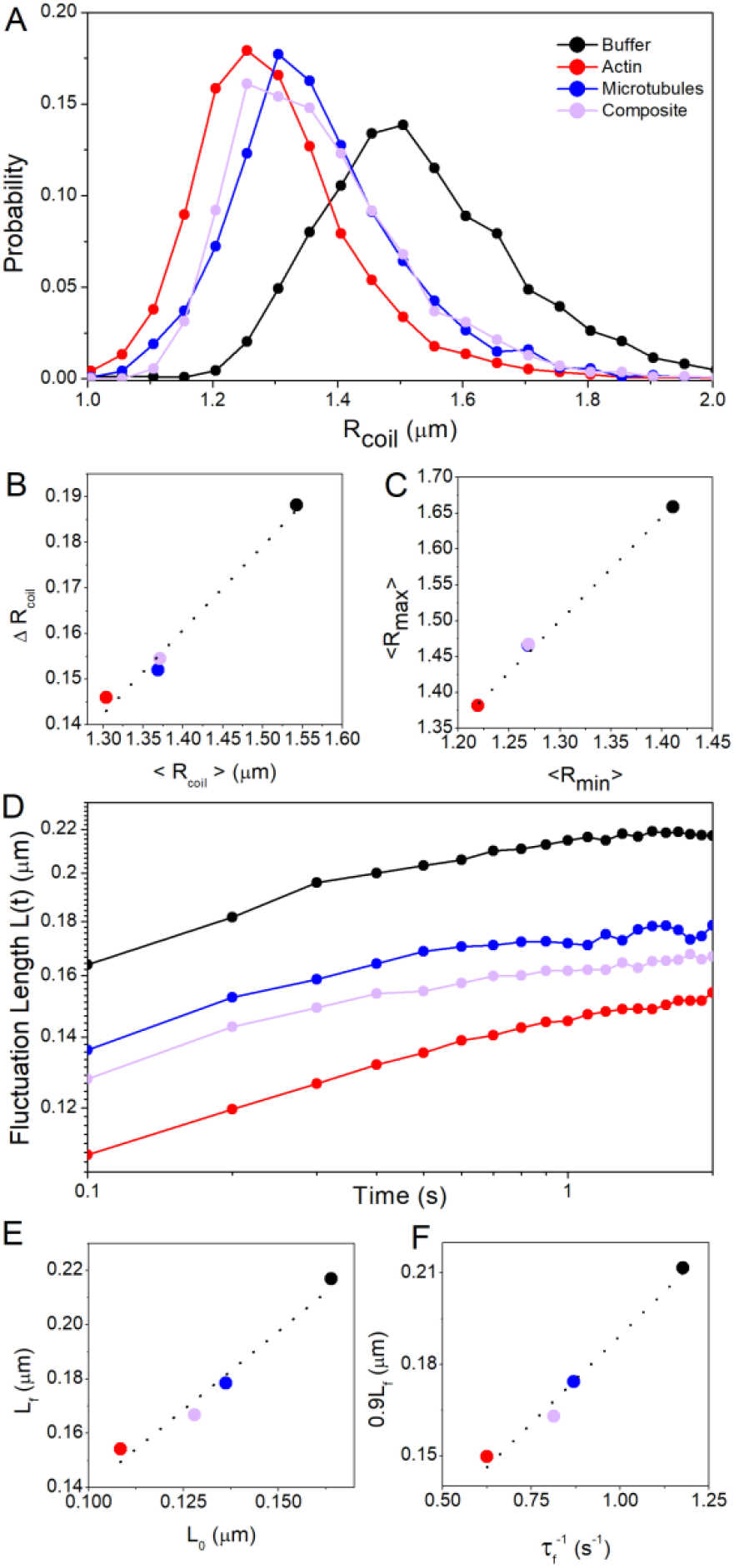
Actin networks induce the most pronounced compaction of DNA and slowing of conformational fluctuations. (A) Probability distributions of DNA coil sizes *R_coil_* show network-dependent compaction of DNA. Distributions for microtubules and composites exhibit significant overlap while actin induces more pronounced compaction (distribution shifted to the left). (B) Standard deviation of coil size distribution *ΔR_coil_* versus mean coil size *<R_coil_>* with dashed line showing the linear relation *ΔR_coil_ = 0.19<R_coil_>*. Actin filaments induce the highest degree of DNA compaction with the least amount of variance while microtubules and composites induce near identical compaction patterns. (C) Mean major axis length *<R_max_>* versus minor axis length *<R_min_>* with dashed line showing the relation *R_max_ = 1.4R_min_*. While the network type determines the degree of compaction, DNA conformations in all cases exhibit similar shape characteristics with eccentricity typical of random coils. (D) Fluctuation length *L(t)* vs time shows that DNA fluctuates between different conformational states with fluctuation rates and lengthscales dependent on crowding conditions. (E) The final fluctuation length *L_f_* vs initial length *L_0_* with dashed line indicating the linear relation *L_f_ = 1.16L_0_*. (F) 90% of *L_f_* vs the rate *τ_f_^-1^* at which *L(t)* reaches 0.9*L_f_* with dashed line showing the linear relation *0.9L_f_ = 0.12τ_f_^-1^*. As shown in (E,F), actin suppresses the lengthscale and rate of fluctuations most appreciably while composites exhibit suppression intermediate between that of actin and microtubule networks.

From the *R_coil_* distributions we evaluate the average size, <*R_coil_*>, as well as the breadth of sizes accessed by the ensemble of molecules over time, quantified by the standard deviation of coil sizes *ΔR_coil_* (Fig. 3B). As shown, *ΔR_coil_* scales proportionally with *R_coil_*, indicating that cytoskeletal crowding does not affect the degree of conformational heterogeneity. However, a clear distinction emerges between actin networks versus microtubule networks and composites. Microtubules and composites exhibit near identical effects on DNA size while actin induces appreciably more compaction. A similar effect is apparent when evaluating the DNA shape or eccentricity, which we quantify by comparing *<R_max_>* to *<R_min_>* (Fig. 3C).*<R_max_>*:*<R_min_>* is 1 for a spherical conformation, ~1.3 for a random coil,^37,38^ and > 1.3 for elongated conformations. As shown, no deviation from random coil shape is evident upon cytoskeletal crowding. However, Fig. 3C corroborates that actin networks induce the most extreme compaction, shown as the smallest *<R_max_>* and *<R_min_>* values, whereas microtubules and composites induce near identical, less extreme, effects on size and shape.

Previous experiments on large DNA crowded by synthetic polymers have also reported molecular compaction arising from the well-known depletion interaction in crowded systems.^6,39,40^ Namely, to maximize the volume available to small crowders, and thus maximize their entropy, large macromolecules are forced to condense to reduce the excluded volume or ‘depletion zone’ that their contours impose. The degree of compaction or volume reduction has been reported to depend on the shape, structure, and concentration of crowders.^6,8,15,17,39,41–46^ While the compaction we measure likely arises from a similar effect, our results are the first, to our knowledge, to report on depletion-driven compaction in highly entangled, viscoelastic systems or when the crowders are comparable in size or larger than the compacted polymer. The varying degree of compaction in different networks that are at equal molarity further indicates the important role that molecular entanglements and stiffness can play in the excluded volume effect.

To determine the time-dependence of conformational states and their dependence on the crowding network, we track the change in *R_max_* over different lag times *t*, termed the fluctuation length *L(t) = <|R_max_(0)-R_max_(t)|>* (Fig. 3D). *L(t)* quantifies both the rate and lengthscale over which DNA molecules fluctuate between different conformational states. As shown, in all conditions *L(t)* increases with time from an initial value *L_0_* to a final, near steady-state value *L_f_*. However, the extent to which *L(t)* increases from *L_0_*, as well as the rate at which it increases, is highly dependent on the type of cytoskeletal network. In line with our compaction results, actin networks result in the smallest *L_0_* and *L_f_* values (Fig. 3E). However, rather than microtubules and composites inducing near identical results, composite crowding induces conformational dynamics in between that of microtubules and actin. To quantify how quickly DNA fluctuates between different conformational states, we calculate the time *τ_f_* needed for *L(t)* to reach 90% of its final value *L_f_, 0.9L_f_* (Fig. 3F). Interestingly, the fluctuation rate *τ_f_^-1^* increases proportionally with increasing *0.9L_f_*, meaning networks that induce more slowing of conformational dynamics also suppress the lengthscale over which molecular conformations fluctuate. Actin induces the most extreme slowing, proportionally suppressing the maximal length of fluctuations and microtubules have the least slowing effect compared to buffer conditions. Composites induce fluctuation rates and lengths intermediate between actin and microtubules.

To couple these intriguing single-molecule conformational dynamics with network-wide DNA transport properties we perform DDM analysis (Fig. 1, Methods). By comparing the image structure function *D(q,t)* for each condition, it is clear that actin networks and composites drastically slow DNA transport, as evidenced by the slow rise in *D(q,t)* compared to buffer conditions and microtubule networks, as well as the lack of any convergence or plateau at long times (Fig. 4A). Similar to the conformational results described above, microtubules have less of an effect on DNA transport, albeit slowing is still evident. However, unlike the single-molecule conformational results, crowding by composites appears to have nearly indistinguishable effects on *D(q,t)* from that of actin, whereas DNA conformations in composites appear to be determined largely by microtubules.

For each spatial frequency *q* we determine the characteristic decay time *τ* of density fluctuations of lengthscale *q*^-1^ (Fig. 4B). As shown, cytoskeletal networks slow transport (i.e. increase *τ*) across all lengthscales. However, actin and composites exhibit much more pronounced slowing compared to microtubules and similar *τ(q)* values across nearly all scales. To further quantify transport, we fit *τ(q)* to the power-law function *τ(q)* = *(Kq^2^)^-1/α^*, where *α* and *K* are the same as in the diffusion equation *MSD = Kt^α^*. Actin and composite networks result in the smallest *K* values, indistinguishable from one another within error (Fig. 4C). Furthermore, both networks also result in highly subdiffusive transport of DNA (*α_actin_* = 0.5, *α_composite_* = 0.66), whereas the deviation from 1 is weaker for microtubules (*α* = 0.8).

**Figure 4.**
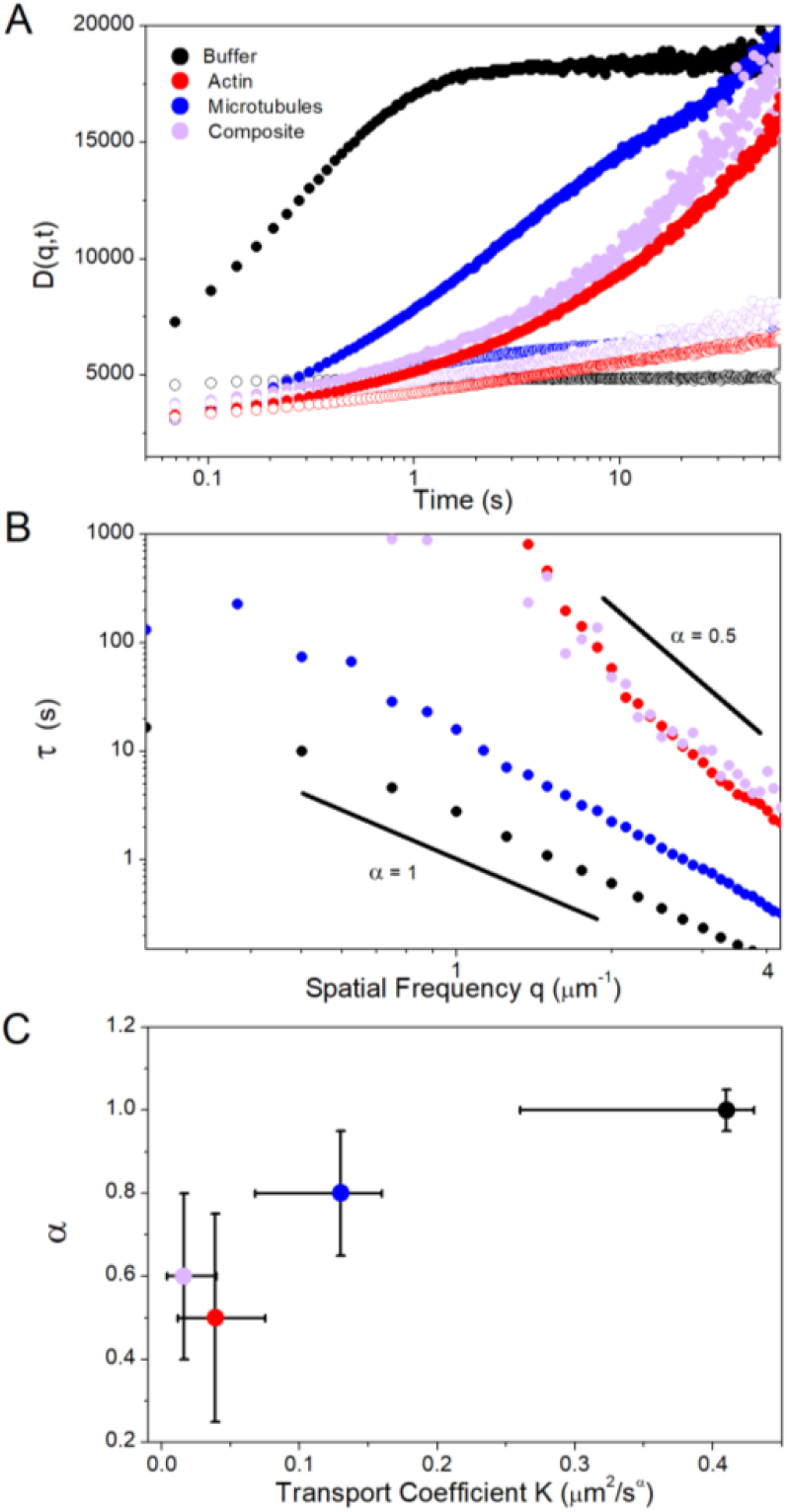
Actin drives extreme slowing of ensemble transport and subdiffusive dynamics of DNA in cytoskeleton networks. (A) DDM image structure functions *D(q,t)* vs time for two different spatial frequencies: *q* = 1.25 μm^-1^ (filled) and *q* = 2.50 μm^-1^ (open). DNA transport exhibits similar slow and anomalous behavior in actin and composites while microtubule networks have a reduced impact on transport. (B) The characteristic decay time *τ* for each spatial frequency *q*, determined via fits to *D(q,t)* (see Methods, Fig. 1). τ(*q*) exhibits power-law behavior that is fit to *τ = (Kq^2^)^-1/α^*. Solid black lines show *α* = 1 (normal diffusion) and *α* = 0.5 (subdiffusion). (C) Transport coefficient *K* versus *α* for each system. Error bars represent the range in *K* and *α* among at least 20 different regions in each of the conditions.

Subdiffusion in crowded systems has been attributed to phenomena such as excluded volume, fractional Brownian motion, heterogeneous environments, viscoelasticity and caging.^4,21,41,47^ Unlike our results, subdiffusion resulting from mobile crowders is often exhibited over a limited temporal range.^21^ This transitional subdiffusion has been reported for small proteins in live cells, and is often attributed to excluded volume and fractional Brownian motion.^21,43^ These studies also found that anomalous transport persisted even with the destruction of the cytoskeletal network, seemingly at odds with our results. Conversely, experiments on small DNA fragments (<3 kbp) in cytoplasm and actin networks showed that the presence of an intact actin network was essential to extreme reductions in the diffusion coefficients of the DNA fragments.^15,16^ However, in these studies normal diffusion (*α* = 1) was reported for all cases.

It is important to note that the proteins and DNA in the described studies were significantly smaller than the DNA in our study. This size difference is of principal importance as the degree of crowding-induced subdiffusion has been shown to depend directly on the ratio of particle or polymer radius to the correlation length of the crowding network.^47,48^ In particular, for colloids diffusing in actin networks, subdiffusion was only apparent when the colloid radius *a* was comparable to the mesh size of the actin network ξ*_A_*, with *α* decreasing linearly with increasing ratios *a*:ξ_*A*_.^47^ In our experiments, the DNA coil radius is *~½R_coil_* ≈ 0.6 – 0.8 μm, whereas the predicted mesh sizes for the actin and microtubule networks are ξ_*A*_ ≈ 0.4 μm and ξ_*M*_ ≈ 0.9 μm, respectively.^49–51^ This comparison suggests that the more extreme subdiffusion in actin networks compared to microtubules may arise from the larger ratio of *R_coil_* to ξ. Further, in the equimolar composite network, ξ_*A*_ is ~2x smaller than ξ_*M*_, such that the actin mesh likely dominates the impact on ensemble DNA transport.^30^

Another critical difference between networks of actin and microtubules is the filament rigidity. Microtubules are ~100x stiffer than actin filaments,^52,53^ and corresponding networks of equal molarity are likewise more rigid and exhibit reduced bending fluctuations and mobility compared to actin networks.^30^ Recent simulations have shown, in fact, that the mobility of large crowders, which results in continuous temporal evolution of the crowding mesh, is essential for true anomalous subdiffusion of trapped polymers.^7^

Subdiffusion that depends on the ratio *a*:ξ has been suggested to arise from caging effects, in which the crowded molecules get trapped in localized ‘pockets’ of the crowded network for extended periods before hopping to new pockets. The distribution of trapping times results in subdiffusion over many decades in time.^4,7,47,48^ Such caging and hopping behavior should also manifest itself as large heterogeneities in transport measured for different molecules in different regions of the network.^4,47^ Within this framework, more extreme subdiffusion (smaller *α*), resulting from more efficient caging, should be coupled with larger heterogeneities in transport. This phenomenon is in fact exactly what we see. Crowding by actin and composites leads to a high degree of heterogeneity in DNA transport compared to microtubule networks. Large-scale spatial heterogeneity is evident in the substantial error bars in *K* and *α* values for these systems.

To explore this heterogeneity phenomenon further, we evaluate the trajectories of single molecules as well as the range of *τ(q)* curves for different regions within each network (Fig. 5). As shown, DNA in buffer conditions exhibits minimal heterogeneities in single molecule transport characteristics, and measurements of *τ* vs. *q* from different regions of the sample all cluster together (Fig. 5A). Microtubule networks exhibit similar consistency among molecular trajectories, and the breadth of *τ(q)* values is not significantly larger than that for buffer conditions (Fig. 5B). Conversely, the distribution of *τ* values measured in different regions of the actin networks and composites spans an order of magnitude across nearly all lengthscales. Also, as evident in temporal color maps, there is a wide range of single-molecule trajectories, from normal Brownian motion (albeit more restricted than in microtubules and buffer) to apparent caging in which the molecule appears nearly stationary over the entire 2 s. This heterogeneity is exactly the reason that computing transport properties using only single-molecule or only ensemble methods becomes so difficult in crowded systems such as these and can lead to inaccurate or incomplete characterization of molecular dynamics.

**Figure 5.**
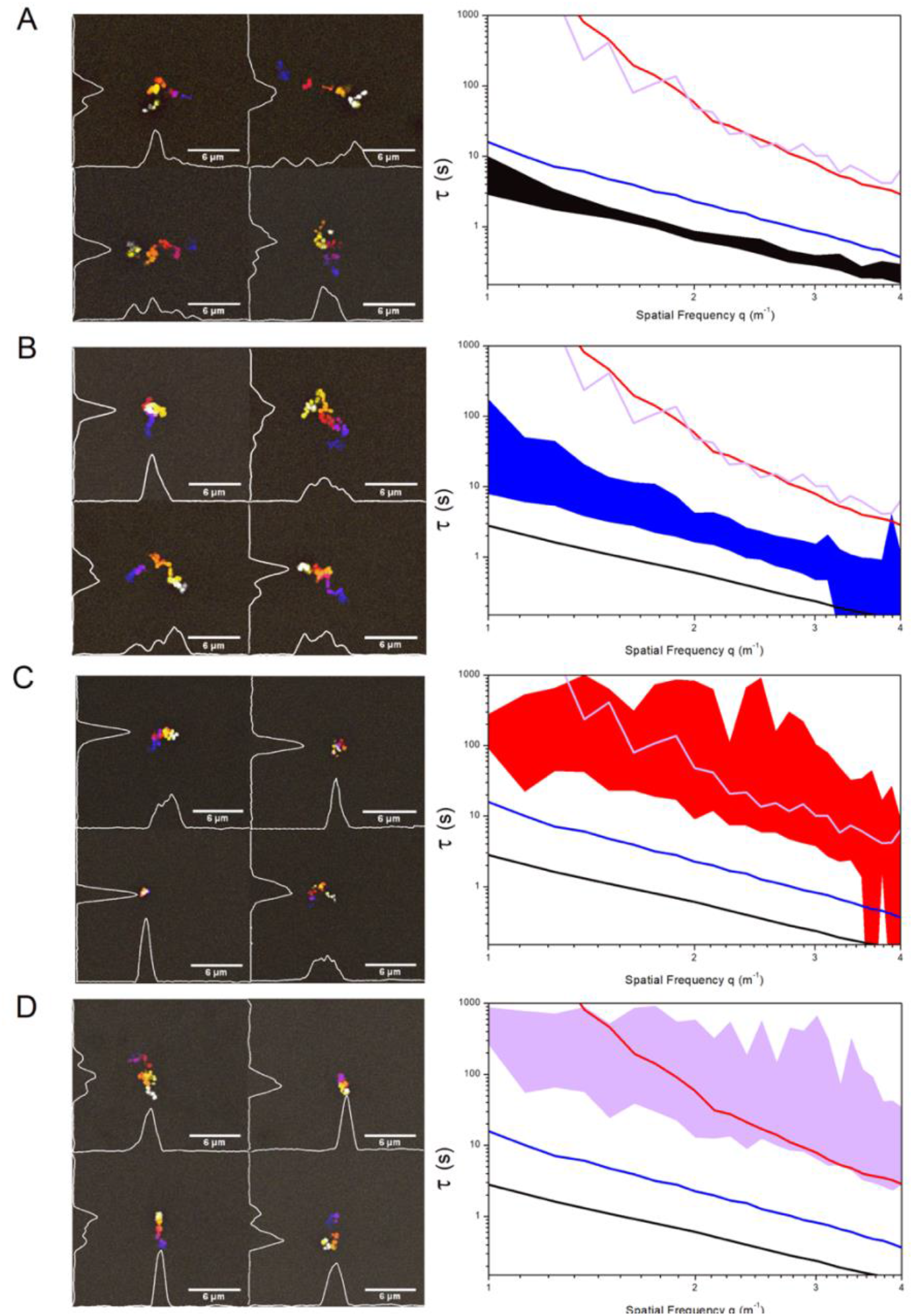
DNA displays heterogeneous transport properties in cytoskeletal networks at both single-molecule and ensemble scales. (Left) Temporal color maps showing sample trajectories of single DNA molecules diffusing in buffer conditions (A), actin (B), microtubules (C) and composites (D). (Right) Characteristic decay times (*τ*) versus spatial frequency (*q*) for each condition. Filled regions indicate the range in data measured in at least 20 different regions of the network. Transport in buffer conditions (A) shows minimal heterogeneity in both single-molecule trajectories and the ensemble distribution of dynamics. Transport in actin and composites (C,D) exhibit a high degree of heterogeneity at both scales. Molecular trajectories are highly varied and the range in ensemble data is quite broad for both systems compared to buffer conditions. Conversely, transport on both scales exhibits much more uniformity, comparable to that in buffer conditions, when crowded by microtubules (B).

## Conclusions

In conclusion, we have coupled recently-developed single-molecule and ensemble techniques (SMCT and SPIDDM) to elucidate the transport properties and conformational dynamics of large DNA molecules crowded by cytoskeletal networks. We span a wide range of spatiotemporal scales (~0.05 – 60 s, ~0.1 – 25 μm) to determine the relationship between single-molecule and network-spanning dynamics, and link DNA dynamics to the structure of the surrounding network. We find that all cytoskeletal networks lead to DNA compaction, suppression of conformational fluctuations, slow and anomalous subdiffusion, and heterogeneous transport. However, the extent to which each of these phenomena occurs depends on the type(s) of filaments in the network. Actin induces the most extreme compaction, fluctuation suppression, transport heterogeneities, and subdiffusion, whereas microtubules have the least impact across all scales.

Composites display a surprising scale-dependent effect, with the presence of microtubules dictating single-molecule conformations and fluctuations, whereas actin filaments drive ensemble transport properties. This complex behavior highlights the importance of characterizing dynamics across wide-ranging scales to paint a complete picture of macromolecular transport in heterogeneous crowded environments. Bridging these scales holds valuable implications for understanding the interplay between intracellular DNA transport and the crowded cytoskeletal environment.

## Methods

### Sample preparation

Double-stranded supercoiled DNA, 115 kbp in length, is prepared *via* replication of bacterial artificial chromosome constructs in *Escherichia coli* followed by extraction and purification as described previously.^54,55^ Supercoiled DNA is converted to linear topology *via* treatment with the restriction enzyme MluI (New England Biolabs).^54^ For microscopy experiments, DNA is labeled with YOYO-I dye (ThermoFisher) at a basepair to dye ratio of 4:1.^54^ Cytoskeleton protein solutions are prepared by resuspending rabbit skeletal actin monomers and/or porcine brain tubulin dimers (Cytoskeleton) in an aqueous buffer consisting of 100mM PIPES, 2mM MgCl_2_, 2mM EGTA, 2mM ATP, 1mM GTP and 5μM Taxol.^30^ ATP and GTP are required to polymerize actin and tubulin into protein filaments (F-actin and microtubules, respectively) and Taxol stabilizes microtubules against depolymerization. 0.05% Tween is added to prevent surface interactions, and glucose (0.9 mg/ml), glucose oxidase (0.86 mg/ml), and catalase (0.14 mg/ml) are added to inhibit photobleaching. Labeled DNA is then added to protein solutions for a final concentration of (1) 0.025 μg/ml for single-molecule tracking experiments (Fig. 1, left) or (2) 20 μg/ml for DDM measurements (Fig 1, right). Final solutions are pipetted into a sample chamber consisting of: (1) a glass slide and coverslip separated by double-sided tape (for SMCT) or (2) capillary tubing that is index-matched to water (for SPIDDM). Chambers are sealed with epoxy and incubated at 37°C for 30 minutes to form an entangled network of actin filaments and/or microtubules (Fig. 2). Networks formed within both types of sample chambers are randomly-oriented and indistinguishable from one another (Fig. 2). Networks consist of either (i) only actin, (ii) only tubulin, or (iii) a 1:1 molar ratio of actin:tubulin, with the total protein concentration fixed at 11.4μM.

### Imaging

For SMCT, DNA is imaged with a Nikon Eclipse A1R epifluorescence microscope with 60x objective and a QImaging CCD camera. Single DNA molecules are imaged and tracked at a frame rate of 10 fps for 20 frames (Fig. 1). All data presented are from an ensemble of ~200 molecules imaged in 2–4 different samples for each condition. Custom-written in-house software (MATLAB) is used to measure and track the major axis length (*R_max_*) and minor axis length (*R_min_*) of each molecule in each frame. We calculate an effective coil size (*R_coil_*) from the major and minor axis length measurements via *R_coil_ = [½(R_max^2^_ + R_min^2^_)]^1/2^* (Fig. 1). Finally, we determine the time-dependence and lengthscale of conformational fluctuations by calculating the fluctuation length *L(t) = <|R_max_(t) – R_max_(0)|>* for a given lag time *t*, which quantifies the lengthscale over which single molecules fluctuate between different conformational states and the timescale over which fluctuations relax to steady-state values. These acquisition and analysis methods, depicted in Fig. 1, have been thoroughly described and validated previously.^3,6,32,56^

For DDM measurements, samples are imaged with a home-built light-sheet microscope with a 10× 0.25 NA excitation objective, 20× 0.5 NA water-dipping imaging objective, and Andor Zyla 4.2 sCMOS camera. Ensembles of DNA molecules are imaged in ROIs of 128 × 1024 pixels (24.2 × 193.5 μm) or 256 × 1024 pixels (48.4 × 193.5 μm) for buffer and protein conditions respectively. Sample videos are taken at 29 fps for 5000 frames (Fig. 1). All data presented in Figures 4 and 5 are from imaging 6 different locations within each sample and 2-4 samples for each condition. For DDM analysis, images are broken into 128 × 128 or 256 × 256 pixel ROIs and analyzed using established DDM routines.^31^ Image structure functions *D(q,t)*, where *q* is the spatial frequency and *t* is the lag time, are fit to a stretched exponential 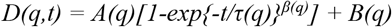 where *A(q)* depends on the optical properties of the sample and microscope and *B(q)* is a background parameter. *τ(q)* values, which represent the characteristic decay time of density fluctuations at each spatial frequency, are determined from fits to *D(q,t)* (Fig. 1), and transport coefficients *K* are then calculated via the relation *τ(q)* = *(Kq^2^)^-1/α^*.^31,33,57^ For a system undergoing normal Brownian diffusion, in which the MSD scales linearly with time, *K* is equivalent to the diffusion coefficient *D* and *α* = 1.

## Data Availability

The data that comprise and support this research is available from the corresponding author upon request.

## Acknowledgements

This work was performed with funding from NIH-NIGMS Award No. R15GM123420 to RMR-A and RM, and AFOSR award no. FA9550-17-1-0249 to RMR-A. The authors would also like to thank R. Dotterweich for her contributions to the initial stages of this project.

## Author Contributions

KR collected data, analyzed data, interpreted data, and wrote the paper. DW collected data, analyzed data, and interpreted data. HR collected and analyzed data. RM designed experiments, analyzed data, interpreted data, and helped write the paper. RMR-A designed experiments, analyzed data, interpreted data, and wrote the paper.

